# A Multi-proxy assessment of the impact of climate change on Late Holocene (4500-3800 BP) Native American villages of the Georgia coast

**DOI:** 10.1101/2021.10.11.463980

**Authors:** Carey J. Garland, Victor D. Thompson, Matthew C. Sanger, Karen Y. Smith, Fred T. Andrus, Nathan R. Lawres, Katharine G. Napora, Carol E. Colaninno, J. Matthew Compton, Sharyn Jones, Carla S. Hadden, Alexander Cherkinsky, Thomas Maddox, Yi-Ting Deng, Isabelle H. Lulewicz, Lindsey Parsons

## Abstract

Circular shell rings along the Atlantic Coast of southeastern North America are the remnants of some of the earliest villages that emerged during the Late Archaic Period (5000 – 3000 BP). Many of these villages, however, were abandoned during the Terminal Late Archaic Period (ca 3800 – 3000 BP). Here, we combine Bayesian chronological modeling with multiple environmental proxies to understand the nature and timing of environmental change associated with the emergence and abandonment of shell ring villages on Sapleo Island, Georgia. Our Bayesian models indicate that Native Americans occupied the three Sapelo shell rings at varying times with some generational overlap. By the end of the complex’s occupation, only Ring III was occupied before abandonment ca. 3845 BP. Ring III also consists of statistically smaller oysters (*Crassostrea virginica*) that people harvested from less saline estuaries compared to earlier occupations. These data, when integrated with recent tree ring analyses, show a clear pattern of environmental instability throughout the period in which the rings were occupied. We argue that as the climate became unstable around 4300 BP, aggregation at shell ring villages provided a way to effectively manage fisheries that are highly sensitive to environmental change. However, with the eventual collapse of oyster fisheries and subsequent rebound in environmental conditions ca. 3800 BP, people dispersed from shell rings, and shifted to non-marine subsistence economies and other types of settlements. This study provides the most comprehensive evidence correlations between large-scale environmental change and societal transformations on the Georgia coast during the Late Archaic period.

## Introduction

The emergence of village life and adaptation to coastal environments are significant transitions in human history that have occurred at various times and places across the globe. Archaeologists in southeastern North America, specifically, have long been interested in social, political, economic, and environmental contexts surrounding the formation and abandonment of early villages along the South Atlantic Coast [1, 2]. Late Archaic Period (5000 – 3000 cal. BP) arcuate and circular shell rings on the Georgia and South Carolina coasts represent what is left of the earliest village communities that emerged in this region. Archaeological research on these circular villages, which predate the adoption of farming, has broadened our understanding of hunter-gatherer economies, the nature of ceremonialism and early monumentality, cooperation, as well as adaptation and resilience in the face of environmental instability [2-4]. However, circular shell ring villages of the South Atlantic coast did not persist across time, and many, especially those of Georgia and South Carolina, were abandoned during the Late Archaic Period. Previous research has focused on the socio-ecological transformations that occurred during the time in which shell ring villages were abandoned in the region, yet few researchers have examined the material record for potential environmental correlations to both the emergence and abandonment of circular shell ring villages. Further, much of the previous research on this topic tends to encompass coarse time scales, lacking the granular resolution necessary to understand how successive generations of people experienced such environmental shifts. Here, we provide a case study from Sapelo Island, Georgia, to document multiple lines of evidence for types of environmental shifts experienced by several generations of villagers that lead to societal transformations on the Georgia coast during the Late Archaic Period.

Circular shell rings along the southeastern Atlantic seaboard of North America emerged around 4400 cal. BP as marsh ecosystems formed in the context of rising relative sea levels, which at that time had reached 1.2 m below present (mbp) [5, 6]. Climatic shifts and relative sea level changes (which may have dropped by as much as 2.5 mbp by 3800 cal. BP and 3.5 mbp by 3100 cal. BP), however, are thought to have led to the eventual abandonment and cessation of shell ring construction in the region [7-9]. Several studies suggest that the abandonment of shell ring villages corresponded with an environmentally correlated collapse in oyster fisheries at this time [7, 8]. Recent research examines the extent to which the hunter-gatherer communities of the Georgia coast underwent reorganization in terms of both settlement and economies to navigate shifting environmental conditions [9, 10]. Specifically, Turck and Thompson [9] argue that hunter-gatherer communities of this region were resilient in the sense that through cooperation and collective agency these communities were able to negotiate shifting social and environmental landscapes in the face of climate change. As climatic shifts changed resource bases (e.g., reduced productivity of oyster reefs], people reorganized their social systems, resulting in changing economies, settlement patterns, and spatial layouts of villages (e.g., the shift to non-shell ring sites that evidence a much-reduced reliance on oysters and other shellfish).

Some of the more well-studied Late Archaic shell-rings villages are located on Sapelo Island, Georgia. Sapelo Island, a barrier island located on the Georgia Coast some 80 km south of present-day Savannah, Georgia USA, plays an important role in our understandings of change and continuity in Native American coastal economies, political organization, and settlement patterns in much of the published literature on the subject over the last two decades (Fig 1). In addition to its archaeological significance, Sapelo Island, and other islands along the Georgia coast, were and continue to be of special cultural significance to Native Americans, such as the Muscogee Nation. The Sapelo Shell Ring complex is located on the northwestern side of Sapelo Island. This site along with research on nearby Ossabaw and St. Catherines islands and regional surveys, has given key insight into the formation and abandonment of villages during the Late Archaic Period [7, 11, 12]. The Sapelo Shell Ring Complex consists of three circular shell rings (Rings I, II, and III) of varying size. Ring I is the largest, consisting of some 5660 m^3^ of shell and covering an area of 6000 m^2^. Previous oxygen isotope analyses (δ^18^O) of mollusk shells and seasonal signatures in archaeofaunal remains from the Sapelo Shell Ring complex indicate that these locales were occupied year-round, with some periods of more intensive gatherings [13, 14]. The Sapelo shell ring villages were likely comprised of coresidential communities characterized by group cooperation and collective action, especially regarding the harvesting of estuarine resources for subsistence and ceremonial purposes [4]. As argued by Thompson [4:30], these villages emerged not because of individuals vying for power and prestige, but rather through the collective agency of groups that worked together to manage dynamic ecosystems that are highly sensitive to human activity and environmental change.

**Fig 1.**
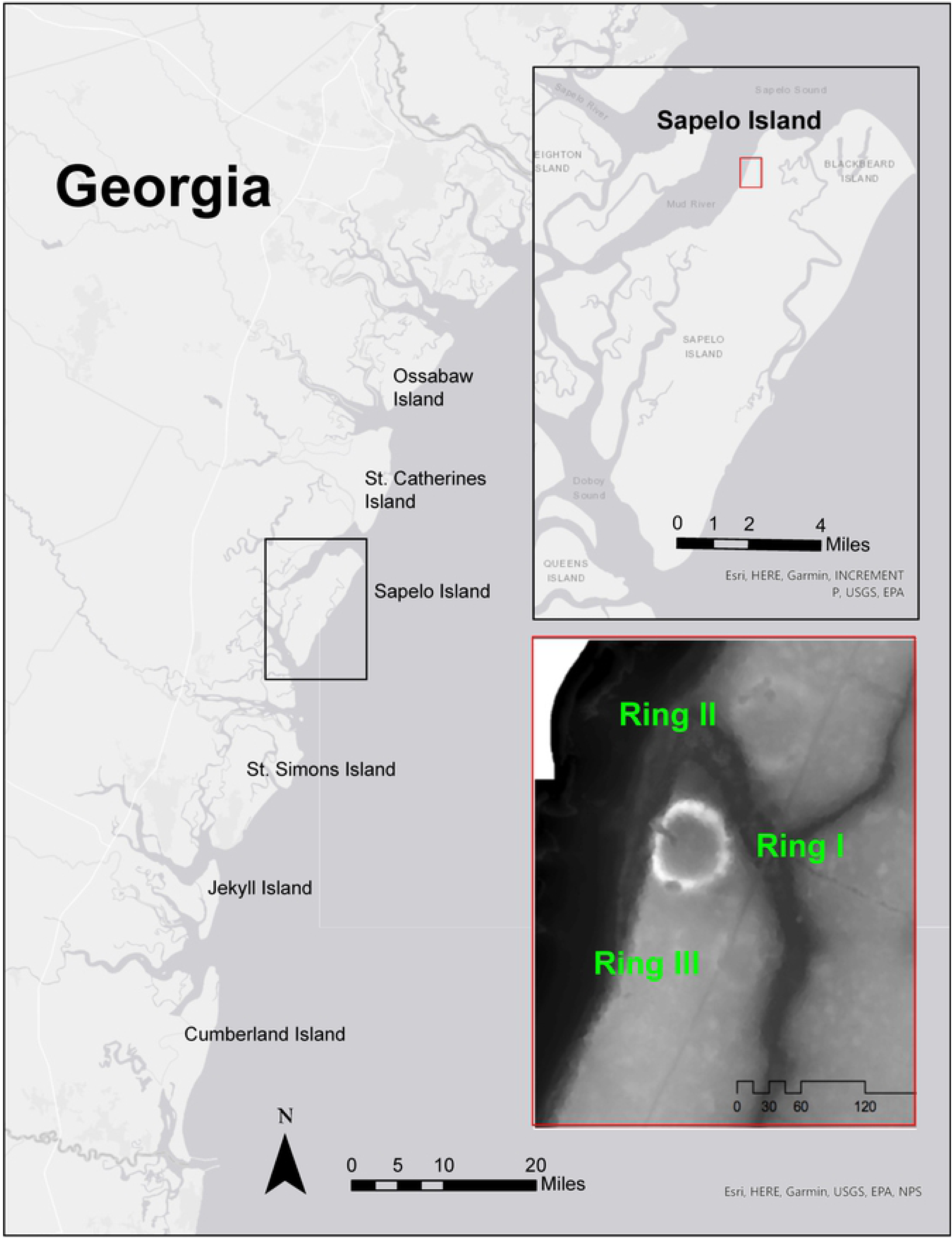
Map of the Georgia coast showing the location of Sapelo Island and shell rings.

As described above, one thing we know for certain is that environmental change played a significant role in the emergence and abandonment of the Sapelo shell rings during the Late Archaic Period. However, exactly what kind of environmental shifts occurred, to what degree, and on what time scale early villagers experienced such shifts have remained elusive. Here, we combine Bayesian chronological modeling of radiocarbon dates with multiple datasets, including oyster morphometrics, stable oxygen isotopes of mollusks, and recent tree ring analyses, to understand the nature and timing of environmental change associated with the emergence and abandonment of circular villages on Sapelo Island, Georgia, during the Late Archaic Period. Our overarching objectives are to: (a) establish a chronological relationship between the three shell rings using Bayesian statistical modeling, and (b) use multiple environmental proxies to document environmental shifts across time that may have led to socio-ecological changes, specifically the formation and eventual cessation of circular shell ring construction on the Georgia coast.

## Materials and Methods

### Radiocarbon Analysis

Establishing a chronological relationship between the three Sapelo shell rings is necessary to link the formation and abandonment of the rings to one another, as well as environmental shifts over time. To examine the chronological relationship of the three rings, we obtained 17 new AMS radiocarbon dates across multiple proveniences from Shell Rings I, II, and III, along with 8 legacy dates. At the request of our Tribal collaborators, we avoided contexts containing ancestral remains in our dating project. Most of our dates come from hickory nut (*Carya* spp.), UID nut fragments, deer bone, pine (*Pinus* spp.), sooted sherds, and UID carbonized wood.

All Accelerator Mass Spectrometry (AMS) radiocarbon measurements were carried out at the Center for Applied Isotope Studies (CAIS) facility at the University of Georgia and followed procedures outlined by Cherkinsky et al. [15]. The charcoal samples were treated following the acid/alkali/acid (AAA) protocol involving three steps: (a) an acid treatment (1N HCl at 80°C for 1 hour) to remove secondary carbonates and acid-soluble compounds; (b) an alkali (NaOH) treatment; and (c) a second acid treatment (HCl) to remove atmospheric CO2. Sample was thoroughly rinsed with deionized water between each step, and the pretreated sample was dried at 105ºC. The dried charcoal was combusted at 900ºC in evacuated/sealed Pyrex ampoule in the present CuO.

The deer bone samples were cleaned by wire brush and washed, using ultrasonic bath. After cleaning, the dried bones were gently crushed to small fragments. The cleaned samples were then reacted under vacuum with 1N HCl to dissolve the bone mineral and release carbon dioxide from bioapatite. The residues were filtered, rinsed with deionized water and under slightly acid condition (pH=3) heated at 80ºC for 6 hours to dissolve collagen and leave humic substances in the precipitate. The collagen solution is then filtered to isolate pure collagen and dried out. The dried collagens were combusted at 575ºC in evacuated/sealed Pyrex ampoule in the present CuO.

The resulting carbon dioxide was cryogenically purified from the other reaction products and catalytically converted to graphite as described in Cherkinsky et al. [15]. Graphite ^14^C/^13^C ratios were measured using the CAIS 0.5 MeV accelerator mass spectrometer. The sample ratios were compared to the ratio measured from the Oxalic Acid I (NBS SRM 4990). The sample ^13^C/^12^C ratios were measured separately using a stable isotope ratio mass spectrometer and expressed as d^13^C with respect to PDB, with an error of less than 0.1‰. The quoted uncalibrated dates have been given in radiocarbon years before 1950 (years BP), using the ^14^C half-life of 5568 years. The error is quoted as one standard deviation and reflects both statistical and experimental errors. The date has been corrected for isotope fractionation.

### Oyster Paleobiology

Eastern oysters (*Crassostrea virginica*), hereby simply referred to as oyster(s), were an important part of larger economic resources on the Georgia coast, and recent research shows that they were sustainably harvested by Native American communities for thousands of years [16, 17]. Oysters were integral to other aspects of life as well, including their use in mound construction and shell ring formation, which can be seen at the Sapelo Shell Rings and later platform mounds along the Georgia coast, such as the Mississippian Period (1150 – 370 cal. BP) Irene Mound [18]. The size of oyster shells is determined by several factors including age, human predation pressures, and environmental variability, with healthier reefs and climatic stability generally producing larger oyster shells [19-22]. For these reasons, temporal changes in oyster size are used as a proxy for environmental change as well as human activity and harvesting practices [21, 22].

To examine if there were any temporal changes in oyster size, we compared the size of eastern oysters between Sapelo Shell Ring I, II, and III. A total of 2,130 eastern oysters were measured from Sapelo Shell Rings I, II, and III. Left valve length (LVL) and left valve height (LVH) measurements (mm) were taken using digital, hand-held calipers, and following a standard method outlined in Lulewicz et al. (22]. All data analyses were conducted using the statistical software R. A Bartlett and Shapiro Wilk test were first used to examine homogeneity of variance and normality of the data, respectively. Since the data are not normally distributed or homoscedastic, a non-parametric Kruskal-Wallis test was used to compare mean LVH and LVL between shell rings, and a post-hoc pairwise Mann-Whitney U test was used to examine which rings are distinguishable regarding mean LVL and LVH. To reduce the possibility of type-I errors associated with multiple comparisons, a Holm correction was used with the Mann-Whitney U test.

### Oyster Geochemistry: Oxygen (δ^18^O] Isotope Analysis

Oxygen Isotope (δ^18^O) analysis of archaeological shell is a widely used method for reconstructing paleoclimate conditions, site occupation histories, and shellfish harvesting practices. Oxygen isotope values in mollusk shells (δ^18^O_carbonate_) are dictated by multiple variables, but largely are a function of the oxygen isotopes composition of ambient water (δ^18^O_water_) [23-27]. Moreover, δ^18^O_water_ covaries with salinity in coastal estuaries [28, 29]. For these reasons, δ^18^O_carbonate_ values in shell can be used to not only trace local environmental changes in the past, but also to explore Native American shellfish harvesting practices, such as season of collection and the range of habitats used for collection [30-39]. Here, we specifically use δ^18^O_carbonate_ values to retrodict the salinity of the habitats where people harvested shell. As with shell size, changes in estimated salinity across time may point to changes in harvesting practices or environmental change. Scholars commonly use two species of mollusks in these studies: hard clams (*Mercenaria* spp.) and eastern oysters, which both have a wide salinity tolerance and are often found in close association. Hard clams tolerate salinity ranges between approximately 17 psu to 37+ psu, with optimal growth between 20 to 30 psu [40, 41]. Oysters’ salinity tolerance is slightly wider, from approximately 5 psu to 37+ psu, with optimal growth conditions between 14 and 28 psu [42, 43].

Incremental oxygen (δ^18^O) isotope analysis was conducted on both eastern oysters (n=19) and hard clams (n=59) collected from all three shell rings. Twenty of these shells were recently sampled; the rest are previously published data from Andrus and Thompson [30]. Laboratory protocols for δ^18^O analysis were adapted from previous studies and are described elsewhere [15, 30, 36]. Briefly, only left oyster valves with a complete chondrophore and clam shells with an intact edge were selected for analysis. Shells with epibiont activity were excluded from analysis as they were likely dead when they were collected [44, 45]. Next, oyster shells were bisected along the chondrophore and clams along their axis of maximum growth. The bisected shells were then mounted onto a slide using Crystalbond™ adhesive and cut into approximately 0.5-inch-thick sections using a slow-speed diamond wafering saw. Each shell was sampled following ontogeny using a Grizzly Benchtop micro-milling system. For oysters, sampling targeted calcitic areas of each shell and avoided aragonite regions [46]. Sampling trajectories followed growth increments as seen in reflected light (in the chondrophore region of oysters and the middle shell layer in the clams) [15]. Generally, 12-20 samples were obtained from each shell, which captured approximately one-year’s worth of growth prior to capture.

The resultant powered carbonate samples were weighed using tin capsules and transferred into Exetainer^®^ 12 ml borosilicate vials. All samples were analyzed for δ^18^O and δ^13^C using a Thermo Gas Bench II coupled with either a Thermo Delta V or Thermo Delta Plus isotope ratio mass spectrometer in continuous flow mode at the University of Georgia’s Center for Applied Isotope Analysis. The values for each sample are reported in parts per mil (‰) relative to the VPDB standard by correcting to multiple NBS-19 analyses (typically 14) per run. NBS-19 was also used to assess and correct for drift and sample size linearity if needed. Salinity values were estimated from shell δ^18^O values following published methods established for the local environments around Sapelo Island [15, 47, 48]. Equations 1 and 2 were first used to estimate δ^18^O_water_ values for each clam and oyster, respectively. The estimated δ^18^O_water_ values were then used to predict salinity for each shell using equation 3. Comparisons of estimated salinity were done between each shell ring for both species combined and each species separately.

### Equations

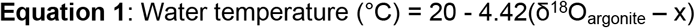

whereas: 31°C is assumed to be the threshold of summer growth cessation for clams (28]; δ^18^O_argonite_ is the most negative value in each clams’ profile; and x = δ^18^O_water_.

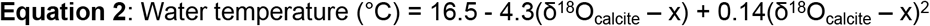

whereas: 28°C is assumed to be the threshold of summer growth cessation for oysters; δ^18^O_argonite_ is the most negative value in each oyster’s profile, and x = δ^18^O_water_. Additionally, a 0.2‰ correction was applied to convert VPDB to VSMOW [47].

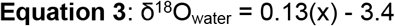

whereas: δ^18^O_water_ is calculated by equation 1 or 2, and x = estimated salinity (psu) [30].

## Results

### Radiocarbon Models

Based on our knowledge of the types of samples, their overall contexts, and stratigraphic ordering, we constructed a series of Bayesian chronological models in OxCal 4.4.4. We then constructed an overall model to determine the ordering of the rings. The structure of the overall model follows closely to the models for each individual ring and can be seen in Fig 2A. We calibrated and modeled all dates using the IntCal20 curve [49], rounding to the nearest 5-year interval [50].

**Fig 2.**
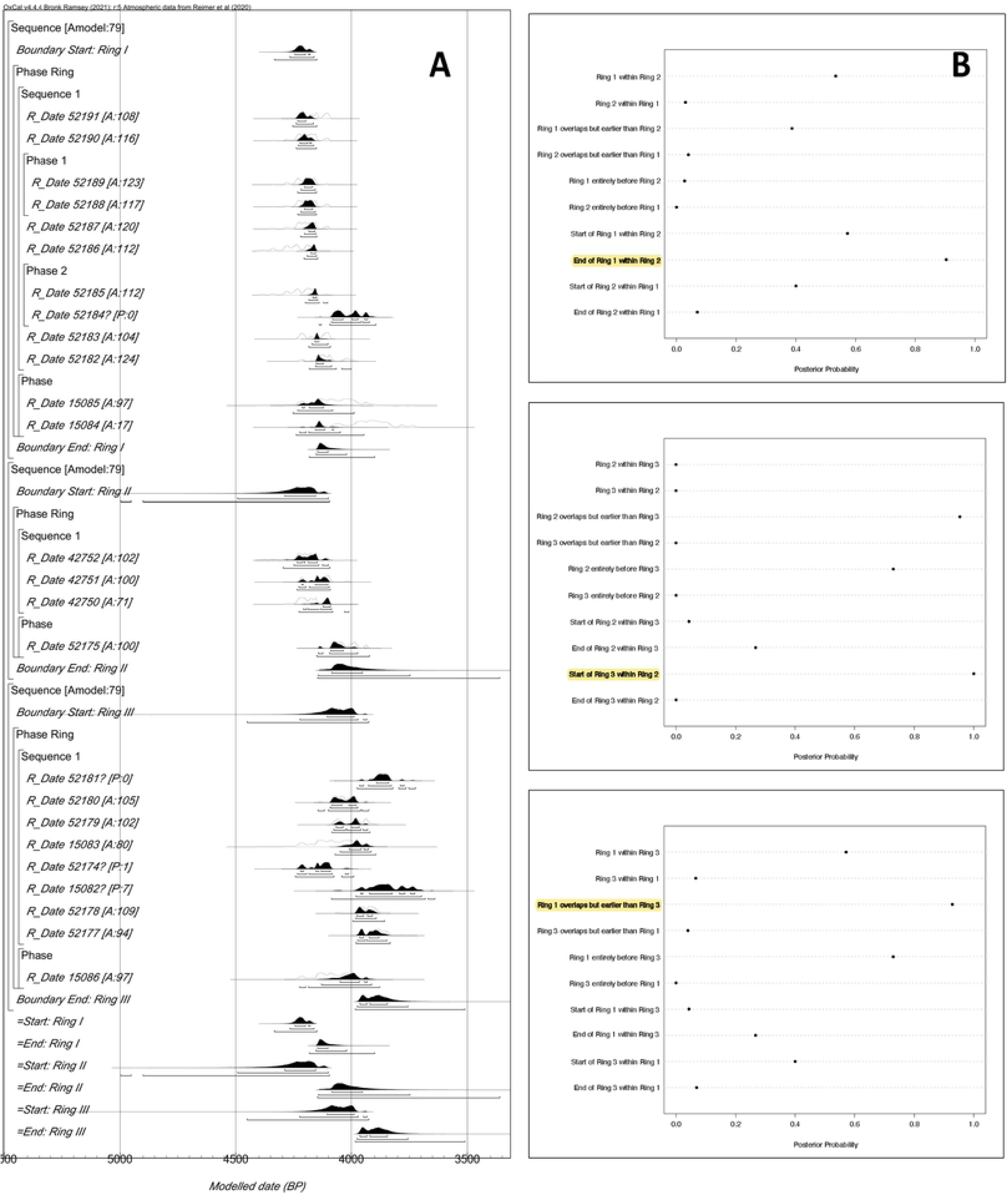
AMS models: (A) Probability distributions; (B) Posterior probability of the chronological relationships for the start and end date of the Sapelo shell rings.

The results of our modeling of the dates indicate good agreement. Both the Amodel (79.2) and Aoverall (79.9) for the model indicate statistical significance, exceeding the 60-threshold established for Bayesian chronological analysis (49, 51]. Due to the long tails in the distribution of these dates, here we focus on the 68% probability range; however, we also provide the 95% ranges (Table 1). All dates indicate good convergence (i.e., >95) save one date (R_Date 15084), which appears to be anomalous and may be the result of bioturbation or some other factor. The model estimates a start date for Ring I of 4245–4175 cal. BP and an end date of 4150–4100 cal. BP; for Ring II, a modeled start date of 4290–4155 cal. BP and end date of 4085–3950 cal. BP; and for Ring III, a modeled start date of 4105–3985 cal. BP and end date of 3965–3845 cal. BP (Fig 2A).

**Table 1.**
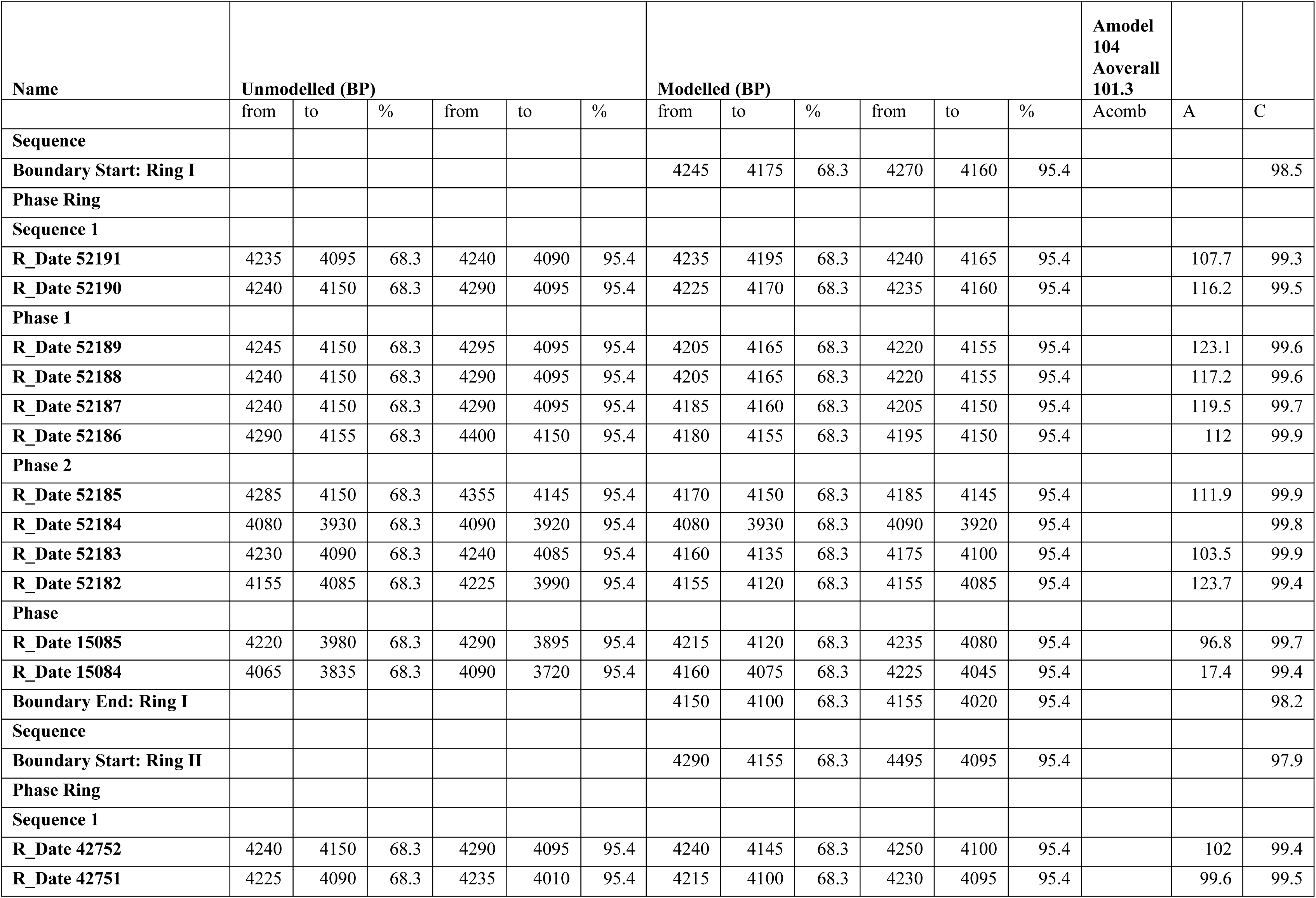
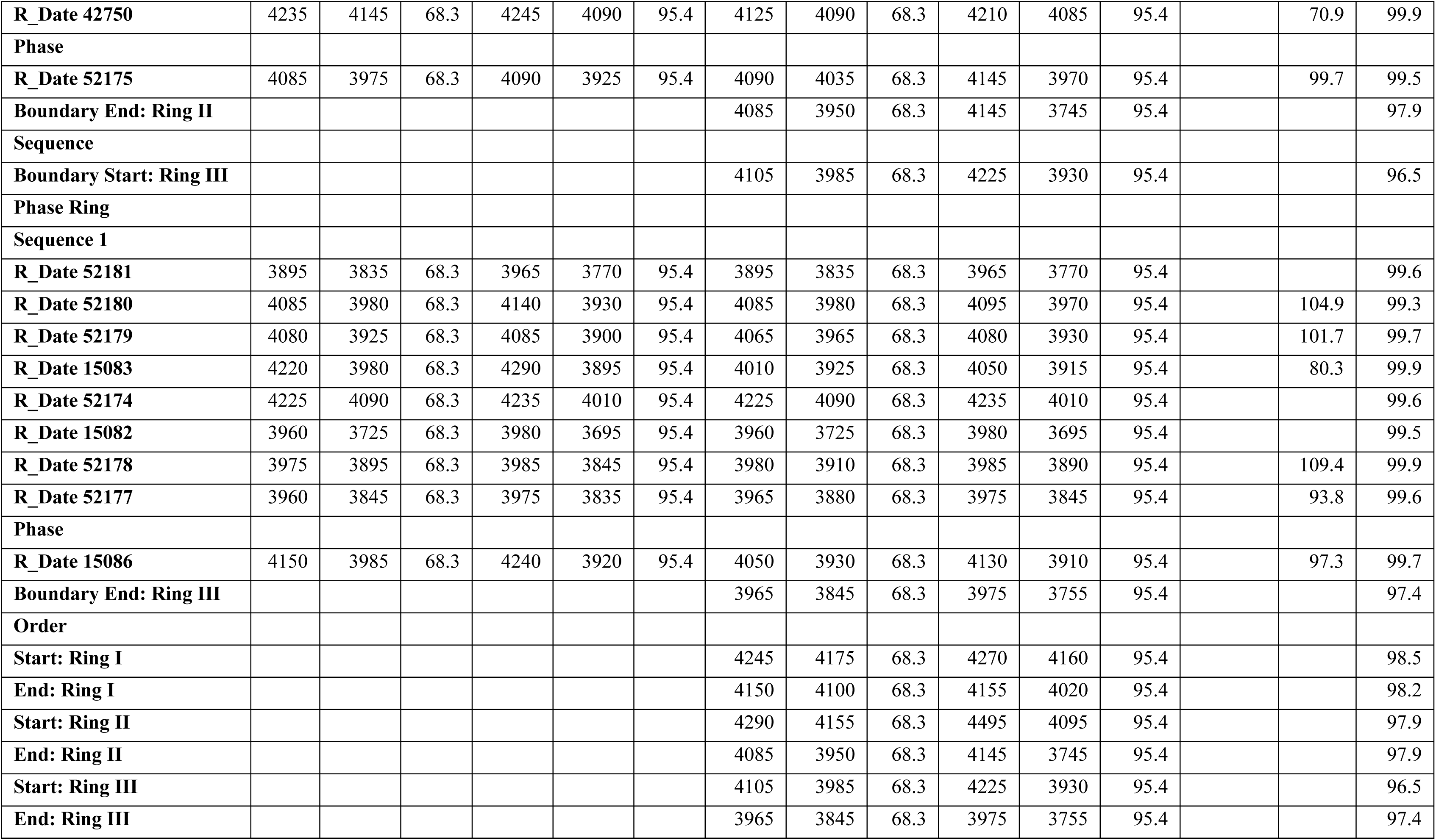
Modeled dates from Sapelo Shell Rings I, II, and III.

To evaluate independently the sequence of occupation of the rings, we used the Order function in OxCal. This function provides probabilities for their relative order based on the dates for each ring. We then used R to calculate the posterior probability for various chronological relationships for the start and end date of the rings on Sapelo (Fig 2B). Based on these results, Ring II appears to be the longest occupied seeing both the rise and abandonment of Ring I. The last generation to occupy Ring II likely also saw the founding of Ring III, which was likely founded after Ring I ceased to be used.

### Eastern Oyster Paleobiology

Our measurements show a clear distinction in oyster size between the three shell rings (Fig 3A, Table 2). Oyster shells from Ring I and Ring II are comparable in size and are generally larger than oysters from Ring III (Table 2). A non-parametric Kruskal-Wallis test indicates that the rings are significantly different regarding mean oyster height (LVH) and mean oyster length (LVL) (LVH: *X*^2^ = 49.5, *p-value* < 0.01; LVL: *X*^2^ = 39.8, *p-value* < 0.01). A post-hoc pairwise Mann-Whitney U test, however, shows that only oysters from Ring III are statistically smaller than Ring I and II regarding both LVH and LVL (at p-value < 0.01). Tests for equality of variance show a significant difference in variation among LVH and LVL between the shell rings, with Ring II exhibiting the greatest variation in oyster size (LVL: p < 0.001; LVH: p < 0.001).

**Fig 3.**
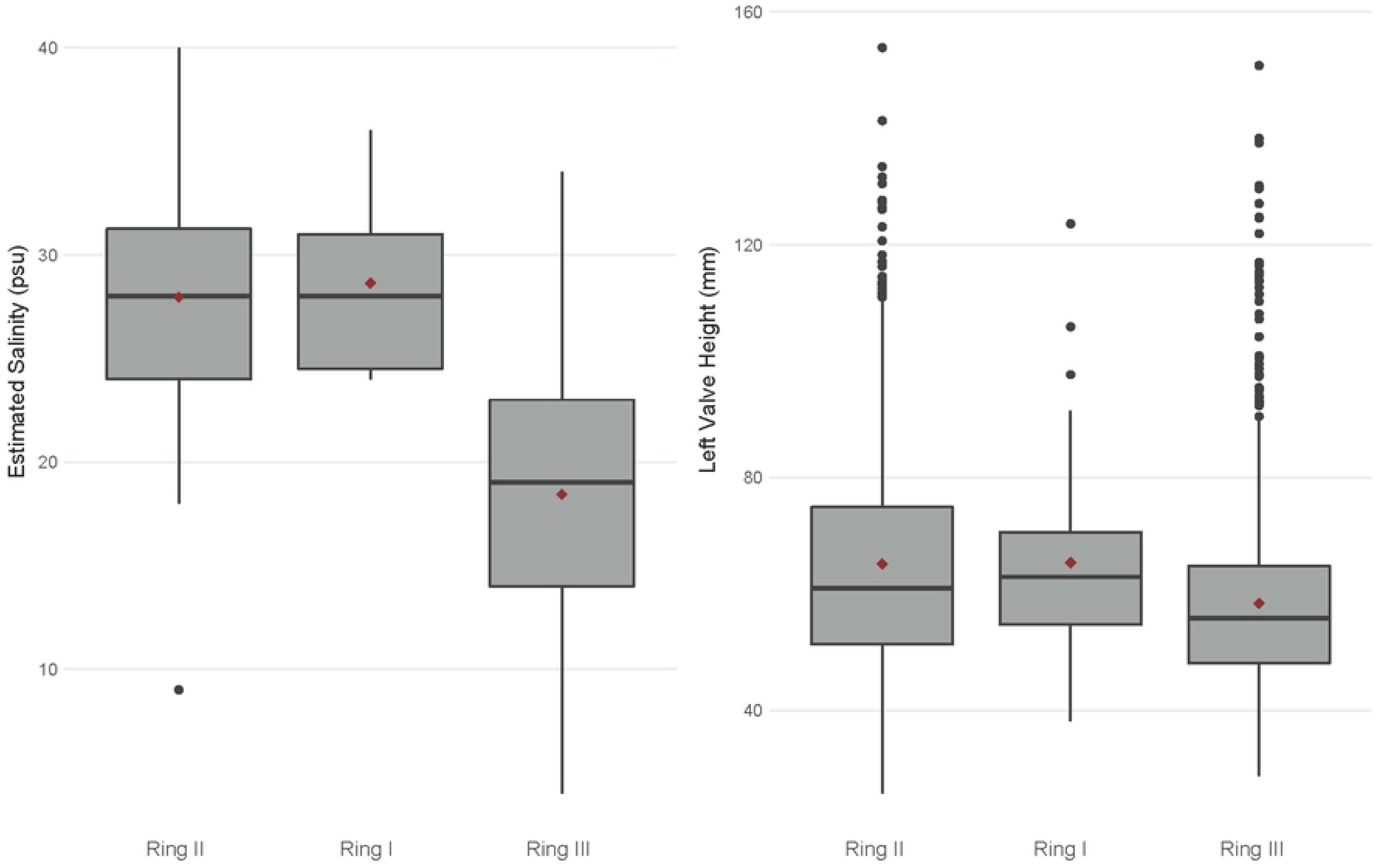
Box plots comparing (A) estimated salinity and (B) mean LVH between the three shell rings, showing significantly lower estimated salinity and smaller shells at Ring III. The shell rings are in chronological order based on the radiocarbon model, and red diamonds show mean values for each ring.

**Table 2.** Descriptive statistics for oyster measurements and oxygen isotope analysis.

| Shell Ring | N (Shell Measurements) | Mean LVL | Mean LVH | N (Shell Isotopes) | Mean $\delta^{18}\text{O}_{\text{water}}$ | Mean Salinity (psu) |
| --- | --- | --- | --- | --- | --- | --- |
| Ring I | 65 | 35.3 | 65.3 | 11 | 0.2 | 28 |
| Ring II | 1057 | 34.1 | 65.3 | 46 | 0.2 | 28 |
| Ring III | 1008 | 32.1 | 58.4 | 21 | -1.1 | 18 |

### Oyster Geochemistry: Oxygen (δ^18^O) Isotope Analysis

Oxygen isotope results show a clear distinction between the shell rings regarding δ^18^O_carbonate_, δ^18^O_water_, and estimated salinity. Oxygen (δ^18^O) values varied among all oyster and clam shells: mean δ^18^O_carbonate_ ranged between −4.0‰ to 0.5‰, and estimated δ^18^O_water_ (using equations 1 and 2) ranged between −2.9‰ and 1.8‰ (Table 3). Most shells also show a general sinusoidal δ^18^O_carbonate_ profile, indicating season fluctuations in water temperature and allowing us to pinpoint summer δ^18^O values (e.g., the most negative value within each shell profile) and predict salinity (Fig 4). Estimated salinity values ranged between 4 and 40 psu, indicating that inhabitants of all three rings were targeting a wide variety of habitats (Table 2). Most estimated salinity values fell within the salinity tolerance for each species, with only three shells falling outside of the expected range. At the mean level, the shell rings are significantly different regarding both δ^18^O_water_ and estimated salinity (δ^18^O_water_: *X*^2^ = 27, *p-value* < 0.01; salinity: *X*^2^ = 32, *p-value* < 0.01). A post-hoc pairwise Mann-Whitney U test, however, indicates that Shell Ring I and II are statistically indistinguishable, and only Ring III is statistically different, with more negative δ^18^O_water_ values and lower estimated salinity (at p-value < 0.01) (Fig 3B). Tests for equality of variance finds that there is no significant difference in variation among δ^18^O_water_ and estimated salinity for each shell ring (δ^18^O_water_: *p-value* < 0.87; salinity: *p-value* < 0.37). These tests remained statistically significant when comparing oysters and clams separately.

**Fig 4.**
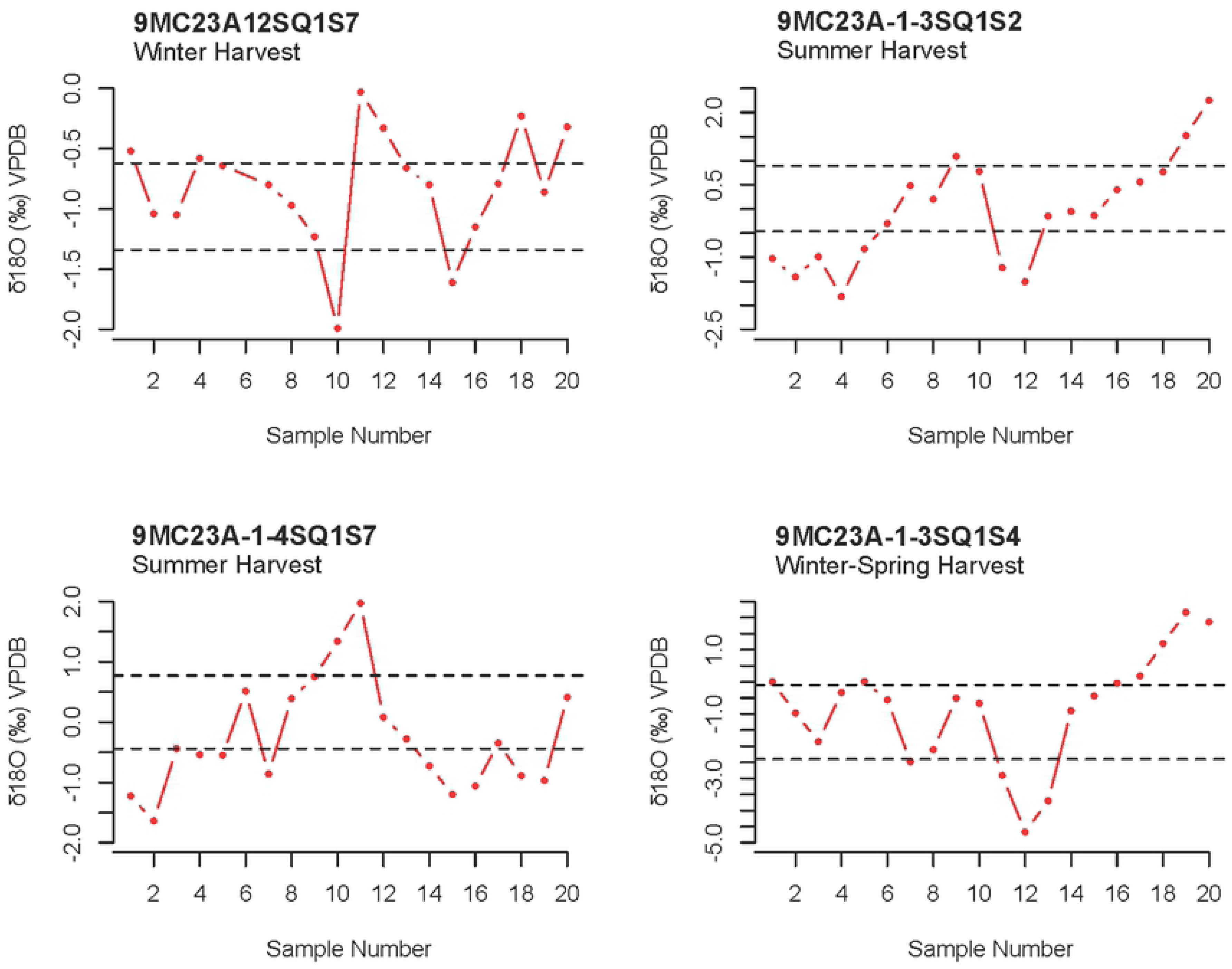
Examples of individual shell δ^18^O_carbonate_ profiles showing seasonal fluctuations in oxygen values and estimated season of harvest. The data sequence follows ontogeny from right to left, with the first value representing time of capture. The dashed lines in each graph represent the values that divide the sample range for each profile into equal thirds (see text above).

**Table 3.** Estimated summer δ^18^O water (**‰** VSMOW) values modeled after Andrus and Thompson’s (2001) oxygen isotope-temperature equations (equations 1 and 2), assuming shell growth cessation at 28°C for oysters and 31°C for clams. Salinity (psu) calculated based on equation 3

| Shell Ring | Species | Sample ID | $\delta^{18}\text{O}_{\text{water}}$ ($\text{‰}$) | Salinity (psu) |
| --- | --- | --- | --- | --- |
| Shell Ring I | *Crassostrea virginica* | OLTS10 | -0.2 | 25 |
| Shell Ring I | *Crassostrea virginica* | OLTS9 | -0.3 | 24 |
| Shell Ring I | *Crassostrea virginica* | OLTS12 | 0.4 | 30 |
| Shell Ring I | *Crassostrea virginica* | OLTS15 | 1.2 | 36 |
| Shell Ring I | *Crassostrea virginica* | OLTS11 | 0.0 | 27 |
| Shell Ring I | *Crassostrea virginica* | OLTS3 | 0.3 | 24 |
| Shell Ring I | *Crassostrea virginica* | OLTS14 | 1.2 | 36 |
| Shell Ring I | *Mercenaria* spp. | CLTS7 | -0.9 | 29 |
| Shell Ring I | *Mercenaria* spp. | CLTS6 | 0.2 | 28 |
| Shell Ring I | *Mercenaria* spp. | CLTS4 | -0.3 | 24 |
| Shell Ring I | *Mercenaria* spp. | CLTS2 | 0.7 | 32 |
| Shell Ring II | *Mercenaria* spp. | 9MC23A-1-3SQ1S1 | 1.0 | 34 |
| Shell Ring II | *Mercenaria* spp. | 9MC23A-1-3SQ1S2 | 0.7 | 32 |
| Shell Ring II | *Mercenaria* spp. | 9MC23A-1-4SQ1S7 | 0.8 | 33 |
| Shell Ring II | *Mercenaria* spp. | 9MC23A-1-3SQ1S6 | 1.8 | 40 |
| Shell Ring II | *Crassostrea virginica* | 9MC23A-1-4SQ1S1 | 0.0 | 26 |
| Shell Ring II | *Mercenaria* spp. | 9MC23A-1-5SQ1S1 | 0.6 | 31 |
| Shell Ring II | *Mercenaria* spp. | 9MC23A-1-3SQ1S5 | 0.9 | 34 |
| Shell Ring II | *Mercenaria* spp. | 9MC23A-1-2SQ1S1 | 1.1 | 35 |
| Shell Ring II | *Mercenaria* spp. | 9MC23A-1-2SQ1S4 | 1.5 | 38 |
| Shell Ring II | *Mercenaria* spp. | 9MC23A-1-2SQ1S7 | 0.5 | 30 |
| Shell Ring II | *Mercenaria* spp. | 9MC23A-1-3SQ1S3 | 0.6 | 31 |
| Shell Ring II | *Mercenaria* spp. | 9MC23A-1-3SQ1S4 | -2.2 | 9 |
| Shell Ring II | *Mercenaria* spp. | 9MC23A-1-3SQ1S7 | 0.7 | 32 |
| Shell Ring II | *Mercenaria* spp. | 9MC23A-1-2SQ1S5 | 0.9 | 34 |
| Shell Ring II | *Mercenaria* spp. | 9MC23A-1-2SQ1S3 | 1.5 | 38 |
| Shell Ring II | *Mercenaria* spp. | 9MC23A-1-2SQ1S2 | 0.9 | 34 |
| Shell Ring II | *Crassostrea virginica* | 9MC23A-1-4SQ1S4 | -0.3 | 24 |
| Shell Ring II | *Mercenaria* spp. | 9MC23A-1-5SQ1S6 | 0.5 | 30 |
| Shell Ring II | *Mercenaria* spp. | 9MC23A-1-5SQ1S4 | 1.8 | 40 |
| Shell Ring II | *Mercenaria* spp. | 9MC23A-1-5SQ1S5 | 1.3 | 36 |
| Shell Ring II | *Mercenaria* spp. | C5A | -0.3 | 24 |
| Shell Ring II | *Mercenaria* spp. | C6A | -1.1 | 18 |
| Shell Ring II | *Mercenaria* spp. | C6A | -0.9 | 19 |
| Shell Ring II | *Mercenaria* spp. | C1A | -0.3 | 24 |
| Shell Ring II | *Mercenaria* spp. | C25A | 0.0 | 27 |
| Shell Ring II | *Mercenaria* spp. | C2A | 0.3 | 29 |
| Shell Ring II | *Mercenaria* spp. | C3A | -0.3 | 24 |
| Shell Ring II | *Mercenaria* spp. | C12A | -0.3 | 24 |
| Shell Ring II | *Mercenaria* spp. | C7A | -0.2 | 25 |
| Shell Ring II | *Mercenaria* spp. | C17A | -0.5 | 22 |
| Shell Ring II | *Mercenaria* spp. | C9A | -0.1 | 26 |
| Shell Ring II | *Mercenaria* spp. | C4A | -0.4 | 24 |
| Shell Ring II | *Mercenaria* spp. | C11A | -0.5 | 23 |
| Shell Ring II | *Mercenaria* spp. | C20A | -0.8 | 20 |
| Shell Ring II | *Mercenaria* spp. | C24A | 0.0 | 27 |
| Shell Ring II | *Mercenaria* spp. | C18A | -0.2 | 25 |
| Shell Ring II | *Mercenaria* spp. | C22A | 0.0 | 26 |
| Shell Ring II | *Mercenaria* spp. | C14A | 0.1 | 27 |
| Shell Ring II | *Mercenaria* spp. | C26A | 0.2 | 28 |
| Shell Ring II | *Mercenaria* spp. | C16A | 0.1 | 27 |
| Shell Ring II | *Mercenaria* spp. | C23A | 0.5 | 31 |
| Shell Ring II | *Mercenaria* spp. | C21A | 0.2 | 28 |
| Shell Ring II | *Mercenaria* spp. | C10A | 0.3 | 29 |
| Shell Ring II | *Mercenaria* spp. | C19A | 0.3 | 29 |
| Shell Ring II | *Mercenaria* spp. | C8A | 0.5 | 30 |
| Shell Ring II | *Mercenaria* spp. | C15A | 0.4 | 29 |
| Shell Ring III | *Crassostrea virginica* | O7 | -2.9 | 4 |
| Shell Ring III | *Crassostrea virginica* | O15 | -1.5 | 15 |
| Shell Ring III | *Crassostrea virginica* | O13 | -1.9 | 12 |
| Shell Ring III | *Crassostrea virginica* | O14 | -0.8 | 20 |
| Shell Ring III | *Crassostrea virginica* | O10 | -0.4 | 23 |
| Shell Ring III | *Crassostrea virginica* | O4 | -0.4 | 23 |
| Shell Ring III | *Crassostrea virginica* | O9 | -1.1 | 18 |
| Shell Ring III | *Crassostrea virginica* | O8 | -0.5 | 23 |
| Shell Ring III | *Crassostrea virginica* | O5 | -0.3 | 24 |
| Shell Ring III | *Crassostrea virginica* | O3 | -0.9 | 34 |
| Shell Ring III | *Mercenaria* spp. | C8 | -2.4 | 8 |
| Shell Ring III | *Mercenaria* spp. | C1 | -1.9 | 12 |
| Shell Ring III | *Mercenaria* spp. | C9 | -1.0 | 19 |
| Shell Ring III | *Mercenaria* spp. | C4 | -1.6 | 14 |
| Shell Ring III | *Mercenaria* spp. | C3 | -1.8 | 13 |
| Shell Ring III | *Mercenaria* spp. | C5 | -1.5 | 15 |
| Shell Ring III | *Mercenaria* spp. | C7 | -0.9 | 19 |
| Shell Ring III | *Mercenaria* spp. | C11 | -0.8 | 20 |
| Shell Ring III | *Mercenaria* spp. | C10 | -0.8 | 20 |
| Shell Ring III | *Mercenaria* spp. | C13 | -0.1 | 26 |
| Shell Ring III | *Mercenaria* spp. | C2 | -0.3 | 24 |

## Discussion

This research provides some of the most comprehensive evidence for environmentally correlated societal transformations on the Georgia coast during the Late Archaic period, specifically regarding the formation and abandonment of circular shell villages on Sapelo Island. Our new chronological research indicates that Native Americans occupied the Sapelo shell rings at varying times with some generational overlap. Ring II had the longest occupational history, spanning from 4290 to 3950 cal. BP. Ring II also witnessed the emergence and abandonment of Ring I (4245 to 4100 cal. BP), as well as the rise of Ring III ca. 4105 cal. BP. By the end of the complex’s occupation, only Ring III was occupied before its eventual abandonment around 3845 cal. BP.

Comparisons regarding oyster paleobiology and isotope geochemistry indicate that villagers at the shell ring complex experienced significant shifts in the environment, especially during the time in which Ring III was occupied. Ring III consists of significantly smaller oyster shells compared to Ring I and II. This suggests a temporal decrease in oyster size given that Ring III is the youngest of the three shell rings. This trend is consistent with other studies on the Georgia coast, such as the Late Archaic Ossabaw Shell Ring, which had the smallest shells in its youngest deposit [22]. Furthermore, even though Rings I and II had similar sized oyster shells that are significantly larger than those from Ring III, Ring II exhibits the greatest variation in oyster shell size, overlapping the range in shell size at the other two rings. This is likely attributed to the long occupational history of Ring II, which temporally overlaps with both Ring I and III. There are two ways to interpret the temporal trend in oyster size, though these drivers are not mutually exclusive and may be attributed to both. First, it is possible that oyster populations experienced harvesting pressures that resulted in smaller shells across time. For example, in heavily predated oyster populations, few individuals will make it to old age due to rapid turnover rates [52, 53]. This results in oyster populations characterized by younger and smaller individuals. Environmental instability that affected local ecosystem productivity also may explain the decrease in oyster size across time. Lower salinity environments from reduced sea levels and periodic river flooding from a wetter climate have been shown to led to high oyster mortality, regular intervals of growth cessation, and thus reduced oyster size [54, 55]. Without other environmental proxies, however, it can be difficult to tease apart whether the observed patterns in oyster paleobiology were attributed to environmental change or human activity.

Results from our isotope geochemistry comparisons further support an interpretation that environmental instability was impacting local ecosystems during the Late Archaic period. Oxygen isotope values in mollusk shells point to a shift toward lower salinity values in the estuaries in which mollusks were harvested, specifically during the time in which Ring III was occupied. This temporal pattern can be attributed to several factors, including previously documented changes in sea levels and local rainfall amounts, which both can impact the amount of freshwater input into local estuaries [5, 6]. It is also possible that villagers who lived at Ring III were targeting mollusks further up estuaries, which are characterized by more freshwater input and thus lower salinity values. However, variation in estimated salinity values indicate that villagers at all three rings were targeting a wide range of habitats. This is corroborated by previously publish data on vertebrate remains from Ring III (see supplemental information), which consists of marine fishes from a variety of habitats that could be captured year-round and using a range of fishing technologies [13]. Moreover, recent research shows that Native American communities along the South Atlantic coast sustainably harvested oysters for thousands of years, evidenced by an increase in oyster size from the Late Archaic through Mississippian periods (5000 – 370 cal. BP) [17]. This stands in contrast to an argument that changes in oyster sizes may reflect unsustainable human management practices. Taking all the evidence into consideration, it is likely that the observed patterns in oyster paleobiology and isotope geochemistry presented here were correlated with environmental fluctuations occurring on decadal or generational time scales.

Contextualizing the observed patterns in oyster paleobiology and isotope geochemistry with new climate data derived from tree ring analysis in the locale, as well as our new radiocarbon model, provides a picture of how these early villagers negotiated climate change that would have ultimately been observable across decades and generations. Recent dendrochronological data indicate a period of environmental instability, including high interannual variability in rainfall patterns, between 4300 and 3800 BP, which began to ameliorate post-3800 BP (Fig 5) [56]. These data contrast with previous research suggesting that environmental instability began around 3800 BP, around the time when people abandoned shell ring villages along the South Atlantic coast. Furthermore, this period of instability overlaps with the chronology of the entire Sapelo Shell Ring Complex, and further contextualizes the changes in oyster paleobiology and estimated salinity of targeted estuaries. Ring I was constructed and occupied during a period of high interannual variability in rainfalls as well as a rapid salinity intrusion event that kills multiple cypress trees, which likely contributed to higher estimated salinity values from oysters at both Rings I and II. Moreover, the time during which Ring III was occupied was overall wetter and had fewer very dry years compared to earlier occupations. A wetter environment leading to more freshwater input into local estuaries, in addition to relative sea level change, both explain the lower estimated salinity seen in oyster shells from Ring III.

**Fig 5.**
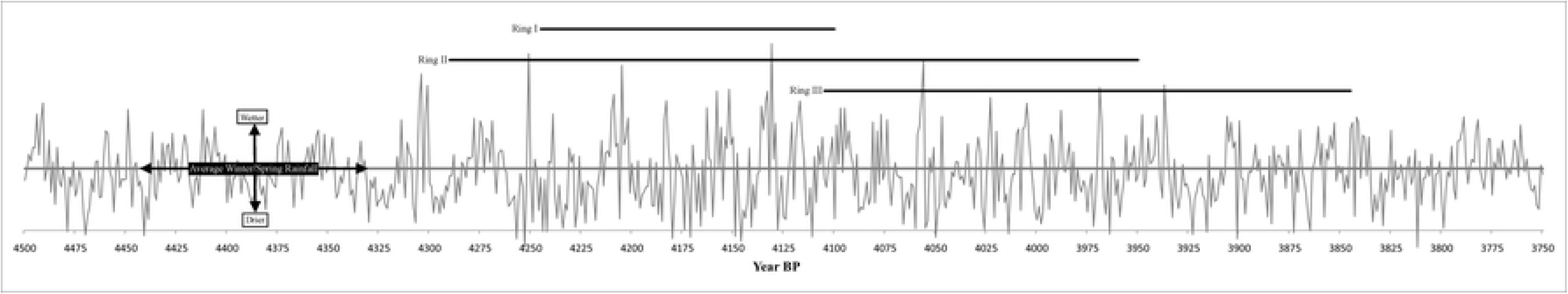
Temporally relevant portion of the multimillennial tree-ring chronology derived from a deposit of ancient buried bald cypress trees at the mouth of the Altamaha River. The chronology is in indices (standardized units representing average ring width, largely indicative in this locale of winter-spring precipitation), with “1000” indicating an annual ring of average width. Enhanced interannual rainfall variability and numerous very dry years are evident beginning around the earliest occupation of Ring I.

These new data shift our focus from not just the abandonment of shell ring villages but also their emergence as an example of resilience in the face of climatic instability. Thompson (2) argues that co-residential aggregation and collective action at the Sapelo shell ring villages would have provided a way to effectively manage oyster and other fisheries that are highly sensitive to environmental change and human activity. Our findings corroborate this argument by providing evidence of climate change and environmental instability experienced by successive generation of villagers on Sapelo Island, leading to societal transformations. As the climate became unstable ca. 4300 BP, Native American communities on Sapelo Island underwent reorganization in both settlement and economies to navigate the shifting environmental conditions. More specifically, through aggregation, cooperation, and collective agency, these communities negotiated changing environmental landscapes in the face of climate change documented here, specifically regarding the management of local fisheries. Given the chronological overlap of the shell rings, knowledge of how to sustainably manage fisheries would have been passed down across generations. Zooarchaeological data from Ring III, as well as the variability in oyster size and estimated salinity values shown here, suggest a persistence of subsistence and fishing strategies characterized by flexibility and use of an array of habitats - a necessity given the daily, seasonal, as well as decadal and generational environmental variability experienced by villagers on Sapelo Island. With continued environmental instability and sea level changes, the construction of shell ring villages ceased, and oyster fisheries on Sapelo Island collapsed ca. 3800 BP. As the climate stabilized post-3800 BP, people in the area shifted to relying on non-marine resources and new settlement patterns for a time (10).

The emergence of village life and adaptation to coastal environments are key transitions in human history that occurred multiple times in a variety of geographic settings. As been the case in other areas of the word (e.g., Peru) where archaeologists intensely study these phenomena, the process by which people became embedded in these landscapes varied widely. Similar to other regions, the Native Americans that established some of North America’s first villages also developed a complexity of ways to adapt to environmental fluctuations and resource shortfalls. This study provides high resolution climate and cultural datasets by which we examine how people reacted to and experienced climate change on a generational level. Climate change is complex and multidimensional as is how people adapt to and mediate their risk in such situations. Our example shows that the emergence of village life among Native Americas created novel social and economic circumstances that revolved around certain estuarine resources (e.g., mollusks). As succeeding generations that occupied the site began to experience climate unpredictability and shifts in resources, occupants decided to alter these patterns to other locations and possible other kinds of social relationships. What is important in this case study is that even though these groups were generationally invested in a specific geographic place, they effectively adapted to changing circumstances and continue to occupy these coastal regions for millennia, albeit in different ways that built on the experience of generations past. This is perhaps a valuable lesson as a host of our current coastal cities and landmarks experience shifting climate and seas.

## Acknowledgements

We thank the Georgia Department of Natural Resources, the Ossabaw Island Foundation, and the Department of Anthropology and Laboratory of Archaeology at the University of Georgia for institutional support. We thank the Historic and Cultural Preservation Department of the Muscogee Nation, especially Raelynn Butler, LeeAnne Wendt, and Turner Hunt for commenting on this manuscript and for allowing us to conduct research on their ancestral lands. This research was supported, in part, in association with the Georgia Coastal Ecosystems LTER project, National Science Foundation grants (NSF Grants OCE-0620959, OCE-123714, 1748276).

## Author Contributions

Designed Research: Garland, Thompson; Performed Research: All authors; Analyzed Data: Garland, Thompson;

Wrote Paper: Garland, Thompson; Edited Final Draft: All authors.

## Data and materials availability

All data needed to evaluate the conclusions are presented in the paper, and all raw will be made available through the Georgia Coastal Ecosystem Long Term Ecological Research Network website: https://gcelter.marsci.uga.edu/

## Supplemental Information Captions

**Table S1**: Sapelo Shell Ring Complex, Ring III, Unit 9 Species List.

**Table S2:** Sapelo Shell Ring Complex, Ring III, Unit 4 Species List.

**Table S3:** Uncorrected AMS dates and context for each sample. See Table 1 for corrected and modeled dates.

**Oxcal Code**

